# Wall following in *Xenopus laevis* is passive

**DOI:** 10.1101/127258

**Authors:** Sara Hänzi, Hans Straka

## Abstract

The tendency of animals to follow boundaries within their environment can serve as a strategy for spatial learning or defence. We examined whether animals of *Xenopus laevis* employ such a strategy by characterizing their swimming behaviour. We also investigated potential developmental changes, the influence of tentacles, which some of the developmental stages possess, and whether wall-following is active (animals seek out wall contact) or passive. Animals’ swimming movements were recorded with a camera from above in a square tank with shallow water and their trajectories were analysed especially for proximity to the nearest wall. With the exception of young larvae, in which wall following was less strong, the vast majority of animals – tadpoles and froglets – spent more time near the wall than what would be expected from the proportion of the area near the wall. The total distance covered was not a confounding factor. Wall following was also not influenced by whether the surrounding of the tank was black or white, illuminated by infrared light, or by the presence or absence of tentacles. Animals were stronger wall followers in smaller tanks. When given a choice in a convex tank to swim straight and leave the wall or turn to follow the wall, the animals consistently left the wall, indicating that wall following in *Xenopus laevis* is passive. This implies that wall following behaviour in *Xenopus* derives from constraints imposed by the environment (or the experimenter) and is unlikely a strategy for spatial learning or safety-seeking.

**Summary statement:** *Xenopus laevis* tadpoles and froglets tend to swim along the walls of a square tank; but this wall following is passive – in a convex tank, they leave the wall.

## Introduction

The exploratory behaviour of animals in unfamiliar environments is often characterized by a tendency to follow walls or distinct borders. Such wall following has been described in mice and rats (Simon et al., 1994; Treit and Fundytus, 1988), where it is often experimentally used as a readout for the level of the animal’s anxiety (Prut and Belzung, 2003; Walsh and Cummins, 1976). Well-studied examples of wall following also include fruit flies and blind cavefish (Besson and Martin, 2005; Goetz and Biesinger, 1985; Liu et al., 2007; Teyke, 1989).

Different potential functions have been ascribed to wall following behaviours. In some cases, wall following might be a defensive strategy; for instance avian predators likely have more difficulties catching e.g. a rat when the latter is moving along a wall compared to when it is moving across an open field (Grossen and Kelley, 1972). This explanation is supported by the fact that rats increase wall following in aversive situations (Grossen and Kelley, 1972), and thus legitimate the use of wall following in rodents as an indicator of anxiety (Gentsch et al., 1987; Simon et al., 1994; Treit and Fundytus, 1988). On the other hand, wall following can also serve as a strategy to learn the spatial setting of an environment. Blind cavefish, which live in dark caves without vision, explore unfamiliar environments by swimming along vertical borders and thereby memorize the layout of the surrounding (Teyke, 1989). A similar spatial learning has also been described in crayfish (Basil and Sandeman, 1999) and humans (Kallai et al., 2005; Kallai et al., 2007), suggesting that wall following is widely used for spatial orientation in vertebrates as well as invertebrates.

However, wall following does not necessarily imply that animals use this behaviour explicitly as a defensive or exploratory strategy. In particular, simply observing a freely moving animal in a concave tank does not indicate whether the animal actively seeks the proximity to the wall. A convex tank, on the other hand, can be used to clearly distinguish between active and passive wall following (Creed and Miller, 1990). The convex curvature allows the animal to chose to either continue straight and leave the wall, or to turn and follow the wall; the latter is then termed active wall following. Accordingly, an animal might appear to be a strong wall follower in a square tank simply because it has no option to make large turns and therefore continues to pursue the border (Creed and Miller, 1990).

To determine whether larvae and adults of the amphibian *Xenopus laevis* tend to swim along the walls of a tank, we quantified the swimming behaviour of these animals in a square concave tank. In addition, tadpole locomotion was recorded in square tanks of different sizes to assess the influence of the size of the environment. Animals at different developmental stages – from small tadpoles (stage 46) to froglets – were employed to estimate the effect of different locomotor styles as well as the role of mechanoreceptive tentacles, which are transiently present at mid-larval stages, in wall following. Finally, a convex tank allowed discriminating between active and passive wall following.

## Materials and methods

### Animals

Experiments were performed on tadpoles and froglets of the South African clawed toad *Xenopus laevis* (*n* = 92) of either sex at developmental stage 46 to 66 (according to Nieuwkoop and Faber, 1956). Stages were identified based on morphological features in freely moving animals in a petri dish under a dissection microscope. All animals were obtained from in-house breeding at the Biocenter of the Ludwig-Maximilians-University Munich, where animals were kept in aerated tanks at 17°C on a 12:12 hour light:dark cycle. All behavioural observations complied with the “Principles of animal care”, publication No. 86-23, revised 1985 of the National Institute of Health. Permission for experiments subjected to approval was granted by the Regierung von Oberbayern (55.2-1-54-2532.3-59-12).

### Image data acquisition – hardware and software

Image data were acquired with two different monochrome cameras from Point Grey (Richmond, Canada; now FLIR Integrated Imaging Solutions) and Point Grey image acquisition software (Fly Capture). The camera was placed in the centre above the tank to record the animal’s movements in the horizontal plane. Videos obtained earlier in the course of the study were acquired using a Grasshopper Firewire camera (GRAS-03K2M-C) with a 640 × 480 resolution at 15 frames per second (fps). These videos were saved as JPG-compressed AVI files. Videos obtained later in the course of the study were acquired using a Grasshopper3 USB camera (GS3-U3-23S6M-C) with a maximum resolution of 1200 × 1200 pixels. The resolution was adjusted depending on animal and tank size and varied from 600 × 600 to 1200 × 1200 pixels with a frame rate of either 15 or 30 fps. Acquired images were saved as LZW-compressed TIFF files. All image data was visually inspected in FIJI (Schindelin et al., 2012; Schindelin et al., 2015), which was also used to create overlays. Further data analysis was performed using Python 3 (Python Software Foundation, https://www.python.org/, see below for details).

### Image data acquisition – experimental conditions

#### Standard procedure

One animal at a time was observed in a 19 × 19 cm Plexiglas tank with a water level of 0.5 to 1.4 cm (0.5 cm only for the smallest animals, otherwise 1.2 - 1.4 cm) at room temperature (20 - 24°C). The vertical walls (20 cm high) of the tank were surrounded on the outer surface by white paper, and the tank was lit from below with four cold light sources placed on either side (ZLED CLS6000, ZETT OPTICS GmbH, Germany) or with a light box (Kaiser slimlite LED, Kaiser Fototechnik, Buchen, Germany) that created an evenly lit area of 46.0 × 20.5 cm. After 1 min adaptation to the environment, a 10 min video sequence was recorded for each of the 92 animals.

#### Tank size

In addition to the recordings of swimming behaviour in the 19 × 19 cm tank, a group of animals (*n* = 9, developmental stages 47 - 50) was also tested in a smaller square tank with floor dimensions of 7 × 7 cm. Animals were filmed for 10 min in each tank; the order of the tank sizes was small first for half of the tested animals, and large first for the other half. All images were acquired with the Point Grey Grasshopper3 camera at 15 fps.

#### Alterations in the illumination

To test for a potential influence of vision, a group of animals (*n* = 10, developmental stages 50 - 65) were filmed successively with both a white and a black paper surrounding the tank, for 10 min each, at a frame rate of 15 fps with the Grasshopper3 camera. The order of black/white was white first for five animals and black first for the other five. A separate group of animals (*n* = 40, developmental stages 53 - 66) was filmed for 10 min both under normal light conditions (see above) and with infrared (IR) illumination (IR Illuminator, TV6700, EcoLine, 850 nm). Because the IR lights also emitted some red light (visible to a human observer), the IR condition most likely was not entirely dark for the animals, but represented a considerably reduced light condition. Half of the animals experienced the normal light condition first, whereas the other half started with IR illumination. Original IR videos lasted 10.5 min and were reduced afterwards to 10 min by removing the first 30 s. The extra 30 s allowed the experimenter to leave the recording room without creating any potentially disturbing light during the 10 min test period.

#### Convex tank

For the analysis of the swimming behaviour of animals in a convex tank, two of the straight walls of the 19 × 19 cm tank were covered with curvatures. Since the number of swimming episodes along the curved walls was limited, image acquisition was manually started and stopped. Occasionally, animals were gently touched at the tail to stimulate swimming towards the convex curvatures and to redirect the swimming trajectory once the animals got arrested in the concave part of the tank. Images were acquired with the Grasshopper3 camera at a frame rate of 15 fps. Unlike the remaining data (see below), videos were not automatically tracked but visually inspected by the experimenter. A ‘trial’ was considered as an animal following the wall and swimming past a convex curve, either following the wall or leaving it. Trials were included independent of the body angles of the animal relative to the wall prior to reaching the curve. Trials were excluded if the animal left the wall before reaching the peak of the convex curve. The remaining trials were scored as ‘going straight’ if the animal departed from the wall at the curve, and as ‘following the wall’ if the animal continued to follow the wall. The proportion of trials in which the animal swam straight was then calculated for all animals with at least 4 trials.

### Tracking of swimming trajectories

Data analysis was carried out by custom-written scripts using Python 3 in the spyder environment (https://github.com/spyder-ide/spyder, version 2.3.8). The main packages included openCV 3 (http://docs.opencv.org/3.0-beta/index.html, version 3.1.0), matplotlib (http://matplotlib.org/, version 1.5.1), numpy (http://www.numpy.org/, version 1.10.4), pandas (http://pandas.pydata.org/, version 0.18.0) and scipy (http://scipy.org/, version 0.17.0). Due to the variety of image file types, image resolution, animal size, illumination conditions and compression quality, the strategy for tracking the animal differed between different sets of experiments. The main difference was that in some cases, background subtraction was carried out before thresholding the image, whereas in other cases images were thresholded directly, either using a simple or a Gaussian threshold.

In contrast, the following steps applied to all cases. The contours of the animals were extracted and the largest contour was taken as the animal. X-Y positions were then calculated relative to the tank geometry. This transformation was achieved by warping the images to the four corners of the tank, which were manually determined. After trajectories were visually inspected, a plot of forward velocity and a video with the animal’s position were generated to ensure that the animal was tracked faithfully. Erroneously tracked frames were identified by visual inspection and spuriously high forward velocities, and their X-Y coordinates were interpolated. Such corrections were necessary in 36 video sequences, 22 of which were animals in the standard condition, with maximally 16 frames to interpolate. In some cases, none of the tracking strategies proved successful, leading to an exclusion of 9 animals in the standard condition.

### Further data analysis

From the X-Y position in the tank-warped images, parameters such as the distance covered during the swimming and the distance to the nearest wall were calculated. To avoid including jitter as animal movement, the trajectories were simplified with the Ramer-Douglas-Peucker algorithm (using the rdp python package, https://github.com/fhirschmann/rdp). The epsilon parameter, which determines the degree of simplification, was set to 10 in a 900 × 900 pixel video, and was scaled linearly to adjust for changes in the resolution. The simplified trajectory was then used to calculate the total distance the animal covered during swimming. Only animals that covered a distance of at least one side length of the tank were included in the analysis; in the standard condition, this led to the exclusion of four animals. A threshold of 15 mm was chosen to define a ‘near wall’ area, and the proportion of time that the animal spent near the wall was calculated. While it is desirable to keep the ‘near wall’ threshold as small as possible, 15 mm was chosen to ensure that the tracked centroid of the large animals was still within that threshold when the animal was near a wall. With 15 mm, the ‘near wall’ area constituted 29.1% of the 19 × 19 cm tank.

When comparing different tank sizes (7 and 19 cm side length), the animals were compared with a 15 mm ‘near wall’ threshold – which might indicate the attractiveness of the wall independent of the size of the tank. However, since the ‘near wall’ area in the 7 × 7 cm tank constitutes 67.3% of the whole tank, the distribution of distances to the wall in both tanks were normalised to the maximum distance, and a threshold was chosen to define the ‘near wall’ area as intermediate in the proportion between the 29.1% and 67.3% that resulted from the 15 mm threshold. Therefore, 0.28 of the maximal distance from the wall was chosen as a threshold for defining the ‘near wall’ area independent of the tank’s size, yielding a ‘near wall’ area of 48% in both tanks, which was intermediate between the ‘near wall’ proportions based on the 15 mm threshold in the two differently sized tanks.

### Code and data availability

The python code used to analyse the data and the tracked data can be found on figshare (Hänzi and Straka, 2017a; Hänzi and Straka, 2017b; Hänzi and Straka, 2017c).

### Statistics and figures

Parameters of interest were tested for normality using a Shapiro-Wilk test; the appropriate parametric or non-parametric tests were chosen accordingly, using an alpha value of 0.05. The distribution of the proportion of time spent near the wall of all animals in the standard condition was not normally distributed; therefore Spearman rank correlations were used to test relationships to other parameters. Figures were assembled in Adobe Illustrator (Adobe Systems Incorporated, San Jose, USA).

## Results

### Swimming trajectories of tadpoles and young adult Xenopus

The swimming behaviour of animals in a square tank between pre-metamorphic stage 47 (larvae) and post-metamorphic stage 66 (froglets) was quantified by monitoring the animals’ trajectories over a period of 10 minutes in each individual (Fig. 1). Examples of animals at different developmental stages revealed a variety of swimming behaviours with respect to the walls of the tank. Independent of developmental stage, some animals exhibited trajectories that appeared to cover the entire tank (Fig. 2A-C), while others swam preferentially along the walls of the tank (e.g. Fig. 2D,G). To visualise the extent of wall following, the cumulative frequency of distances to the nearest wall over the 10 minutes period of swimming was plotted (see Fig. 1B). This graphical presentation is equivalent to a histogram of distances to the nearest wall that are summed up along the X-axis.

**Figure 1.**
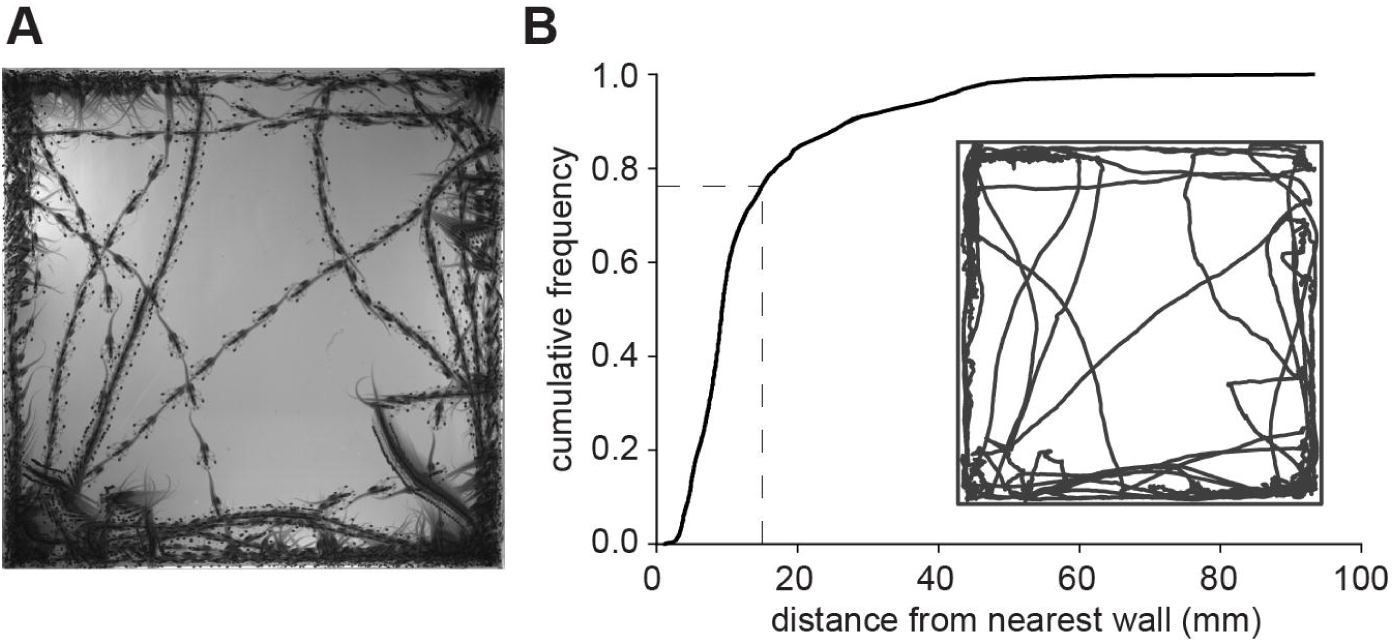
Example swimming trajectory and cumulative frequency distribution of a *Xenopus* tadpole’s distance to the nearest wall. (A) Minimum intensity projection showing the entire trajectory of a stage 54 tadpole (body length 3.6 cm) during swimming in a 19 × 19 cm tank over a 10 min period at a temporal resolution of 3 fps. (B) Cumulative frequency distribution of the animal’s distance to the nearest wall; note that the animal spent over 75% of the time within 15 mm of the nearest wall (dashed lines); the inset shows the tracked trajectory.

**Figure 2.**
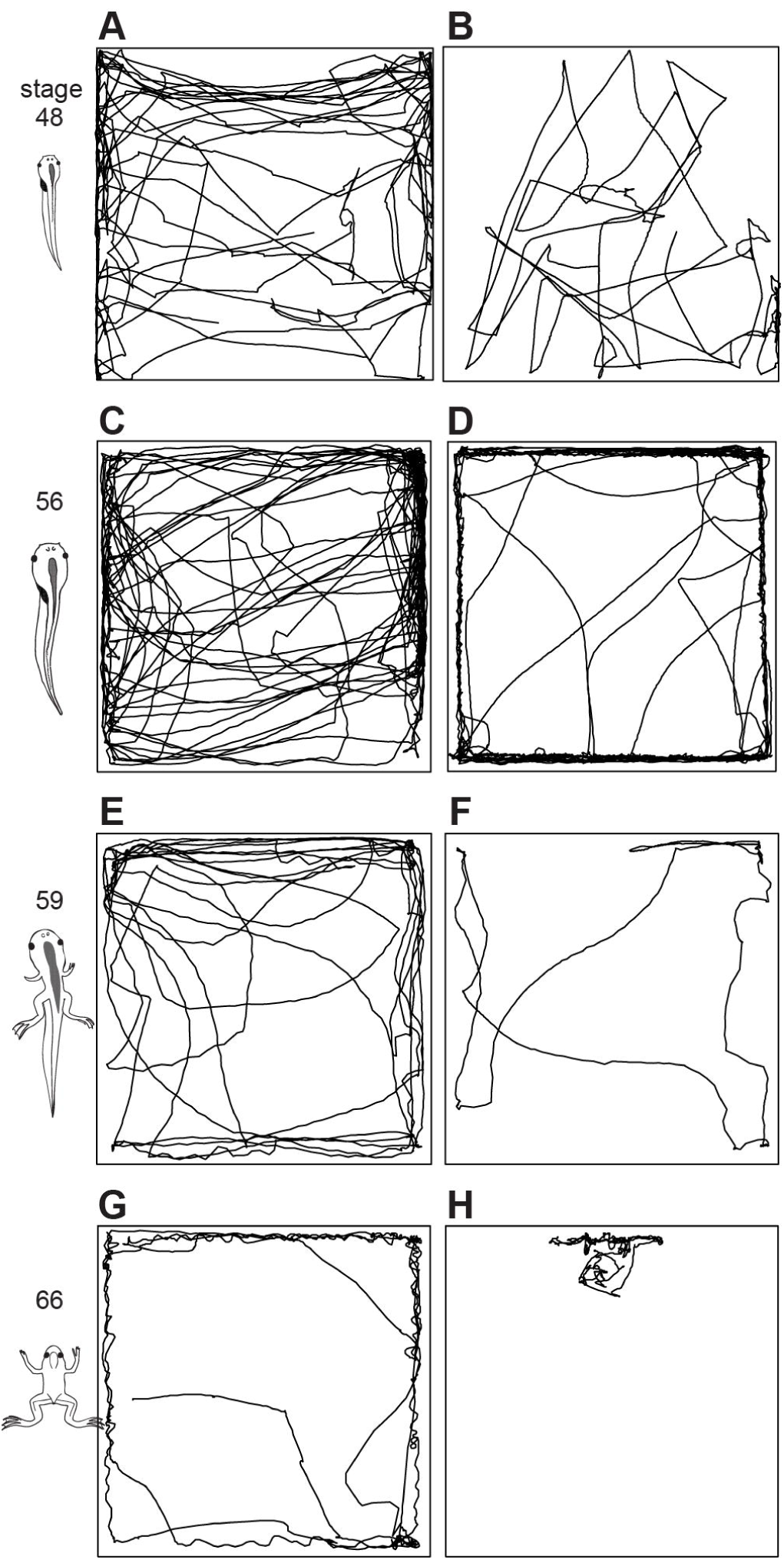
Example swimming trajectories of larval and adult *Xenopus* at different developmental stages. (A-H) Reconstructed trajectories during swimming in a 19 × 19 cm tank over a 10 min period of two animals, respectively, at stage 48 (A,B), stage 56 (C,D), stage 59 (E,F) and of two froglets at stage 66 (G,H). Note the variability of the trajectories of animals at the same developmental stage. The size of the animal schemes on the left (from Hänzi and Straka, 2016) is not related to the spatial dimensions of the trajectories.

The cumulative frequencies of distances to the nearest wall for all animals (*n* = 79) are shown in Figure 3A. The proportion of time that the animals spent near the wall (within 15 mm of the wall) was taken as a measure of the strength of wall following. As a group, the 79 animals differed significantly from the proportion that could be expected from the ‘near wall’ area (29%, Fig. 3B, Wilcoxon signed rank test, *p* < 0.0001). Five animals, however, spent less than 29% of their time near the wall, which is the proportion of the ‘near wall’ area. Four of these were of developmental stage 48 or below and this tied in well with the impression that the strength of wall following increased with developmental stage (Fig. 3C, Spearman’s rank correlation between stage and proportion near the wall, rho = 0.48, *p* < 0.0001, *n* = 79), suggesting that *Xenopus* larvae/froglets become stronger wall followers during ontogeny.

**Figure 3.**
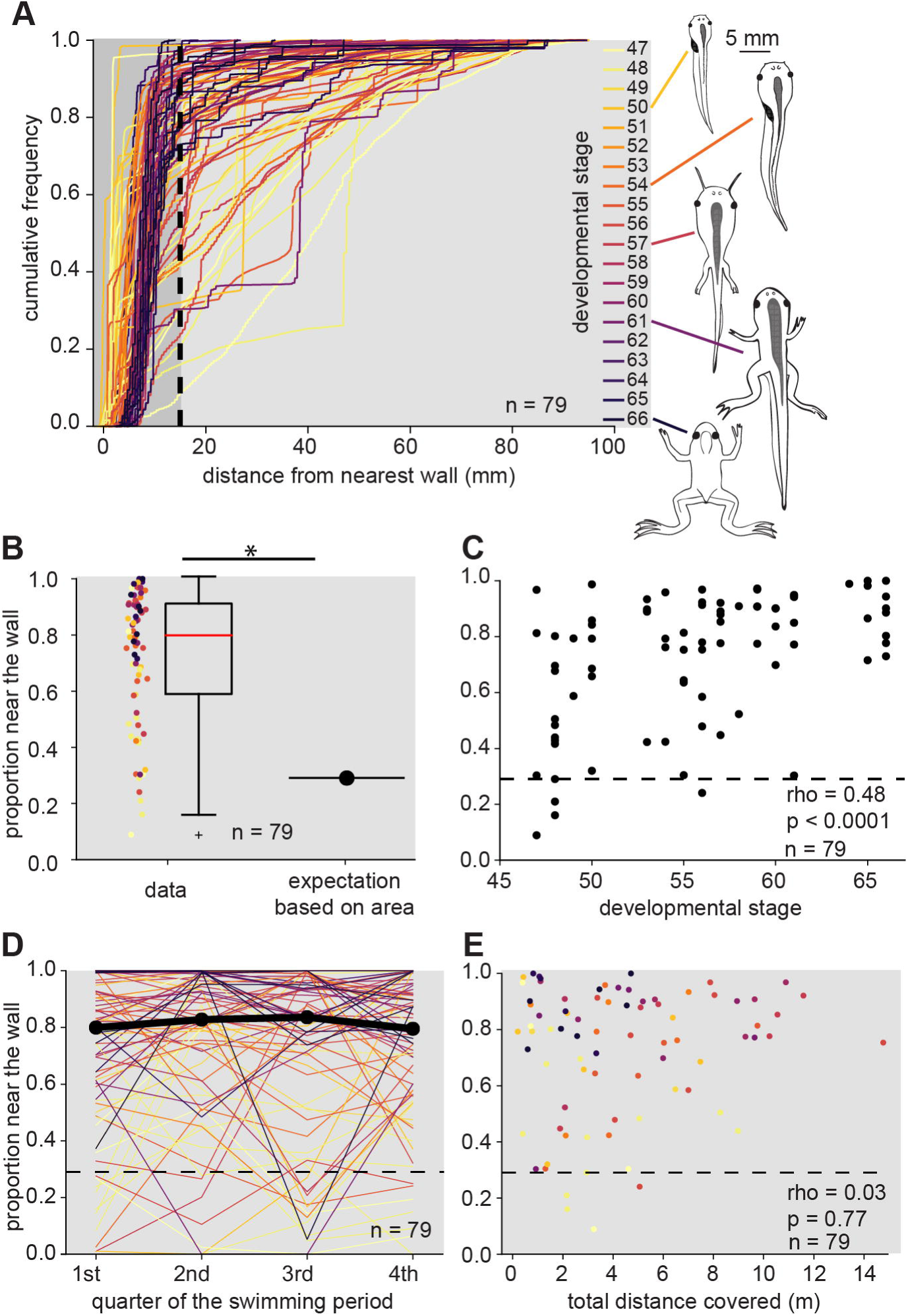
Characterisation of wall following of larval and young adult *Xenopus* during swimming in a square tank. (A) Cumulative frequency distributions of the distance to the nearest wall during swimming of tadpoles and froglets (*n* = 79) for 10 min in a 19 × 19 cm tank; traces are colour-coded with respect to developmental stage (colour-code on the right); dashed black line indicates the threshold of the ‘near wall’ area (15 mm). (B) Proportion of time that the animals spent near the wall from the data shown in A, as colour-coded dots and as a boxplot. The expectation of how much time the animals would spend near the wall based on the ‘near wall’ area as a proportion of the total area is shown on the right. The animals’ proportions were significantly different from this expectation (Wilcoxon signed rank test, *p* < 0.0001, *n* = 79). (C) Relationship between proportion of time that the animals spent near the wall and the developmental stage of the tested animals (*n* = 79); note the significant Spearman’s rank correlation between stage and one-sample KS statistics (*n* = 79, *rho* = 0.48, *p* < 0.0001), indicating that older animals are stronger wall followers. (D) Separate proportion of the time that the animal spent near the wall for each quarter of the 10 min swimming episode shown in A *(n* = 79, colour-coded for developmental stage). The median across all animals is shown as a thick black line. These proportions did not change significantly across the four quarters of the 10 min swimming period (Friedman test, *p* = 0.29). (E) Relationship between the proportion the animals spent near the wall and the total distance covered by an animal over the 10 min swimming period (colour-coded for developmental stage). The absence of significance (Spearman’s rank correlation, *rho* = 0.03, *p* = 0.77) indicates that total covered distance is not a confounding factor for the degree of wall following as measured by the proportion of the time spent near the wall. The dashed line in C-E indicates the ‘near wall’ area as a proportion of the total tank area. Schemes of *Xenopus* in A from (Hänzi and Straka, 2016).

To reveal potential changes in wall following behaviour in individual animals over the 10 minute test period, the respective proportions of time spent near the wall were separately calculated for the four quarters of the swimming period (Fig. 3D). Since the proportions of the four quarters were not significantly different from each other (Fig. 3B, Friedman test, *p* = 0.29), the individual wall following strategy of a particular animal persisted over the entire test period. Moreover, the total distance covered within the 10 minutes was no confounding factor for wall following, since the rank correlation between the total length of the trajectory and the proportion of time spent near the wall was not significant (Fig. 3E, Spearman’s rank correlation, *rho* = 0.03, *p* = 0.77).

### Role of tentacles in wall following behaviour

During larval development between stage 51 and 60, *Xenopus laevis* tadpoles transiently posses a mobile pair of rod-like appendages that protrude from the corners of their mouths (Nieuwkoop and Faber, 1956). These appendages might be necessary or at least advantageous for wall following, given the presence of Merkel cells, potentially assigning a tactile function to these tentacles (Ovalle, 1979; Ovalle et al., 1998). However, contrasting with normal development, a number of animals from our breeding facility failed to naturally develop noticeable tentacles. This allowed to directly test the influence of tentacles on the degree of wall following. Accordingly, the swimming behaviour of a population of tadpoles at developmental stages 54 - 60 without appendages (*n* = 11) was compared with that of an age-matched group of tadpoles (*n* = 13) that possessed tentacles with a length of at least 3 mm.

Statistical analysis of the swimming behaviour as reported above indicated that both populations of animals had a similar propensity for wall following (blue and red traces in Fig. 4A). This is demonstrated by the overlapping distributions of the cumulative frequencies of distances to the nearest wall in animals with and without tentacles (blue and red traces in Fig. 4A). The proportions of time that these animals spent near the wall were not significantly different between animals with and without tentacles (Fig. 4B, Mann-Whitney-*U* test, *p* = 0.09). If anything, animals without tentacles were located closer to the wall than animals with tentacles (see blue and red traces in inset in Fig. 4A). This likely derives from the fact that the presence of tentacles creates an additional distance of the tadpole with respect to the wall that is not present in animals without tentacles. Tentacles are therefore no prerequisite for wall following. This, however, does not exclude that tentacles are used as tactile probes; rather it shows that despite the absence of tentacles, tadpoles follow the walls of a tank and potentially use facial skin areas as tactile probes.

**Figure 4.**
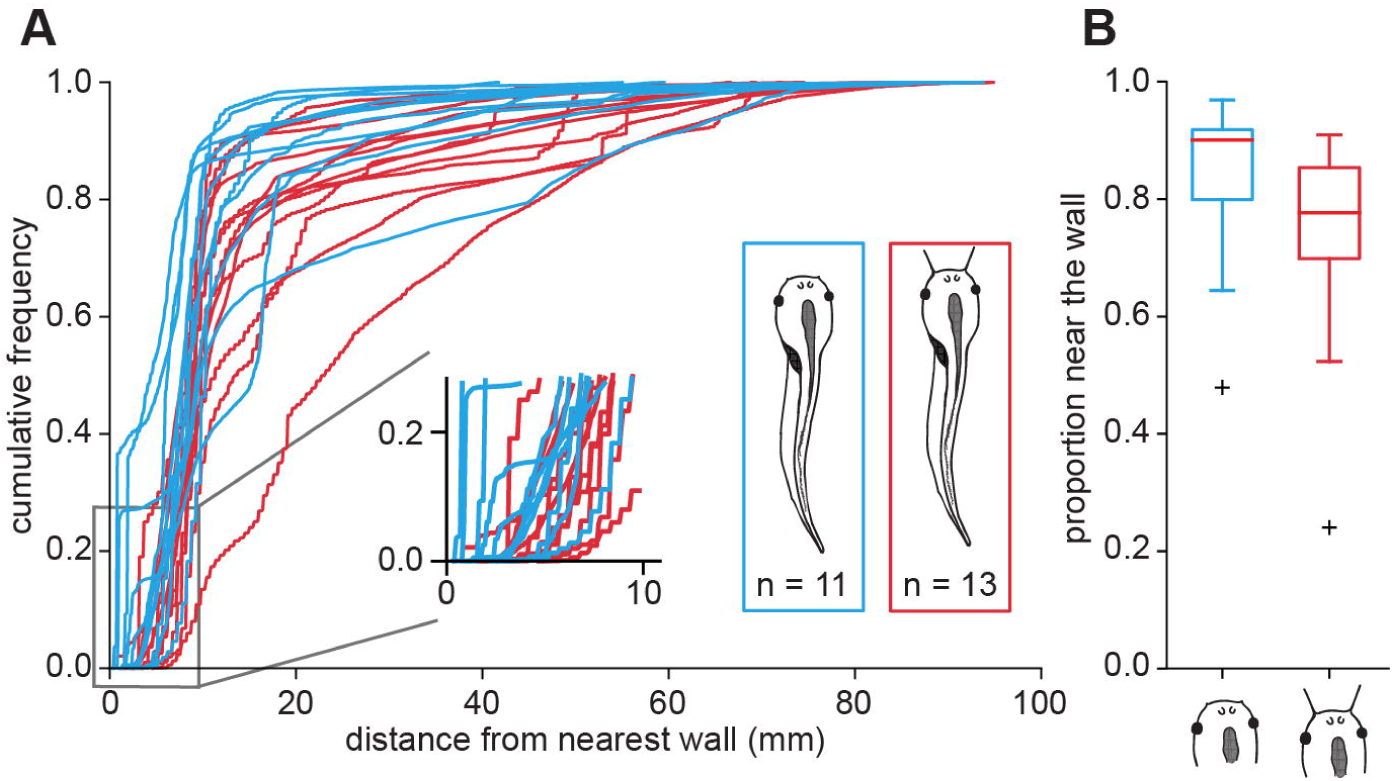
Influence of tentacles on wall following during swimming in *Xenopus* larvae. (A) Cumulative frequency distributions of the distance to the nearest wall of animals with tentacles (red, *n* = 13) and of animals without tentacles (blue, *n* = 11) between developmental stages 54 – 60; the inset is a higher magnification of the initial part of the cumulative frequency distribution and shows that tadpoles without tentacles (blue) align closer with the wall compared to tadpoles with tentacles (red). (B) Proportion of the time that the animals with and without tentacles spent near the wall; the two groups were not significantly different (Mann-Whitney U test, *p* = 0.09).

### Wall following under different luminance conditions

The wall following of *Xenopus* larvae/froglets analysed above was further examined during swimming under different illumination conditions, which could have facilitated or impaired wall detection. A potential influence of the visual system was therefore evaluated in a separate set of experiments where the swimming of stage 50 - 65 tadpoles/froglets (*n* = 10) was compared in a tank in which the four walls were covered on the outside by a white or a black background (Fig. 5A,B). Analysis of the swimming behaviour indicated that the propensity for wall following was not related to the background (Fig. 5B) based on the proportions of time that each animal spent near the wall in the two conditions (paired t-test, *p* = 0.59). This suggests that the visual system exerts no apparent influence on the tendency of *Xenopus* for wall following. This conclusion was confirmed by another set of experiments in which the swimming behaviour of tadpoles/froglets (*n* = 30, stage 53-66) was tested under both white light (cold light source) and infrared illumination (850 nm, Fig. 5C,D). Analysis of the proportion of time spent near the wall revealed no significant difference between the two conditions (Fig. 5D, paired Wilcoxon signed-rank test, *p* = 0.47), indicating that the reduced light condition during infrared illumination had no effect on wall following.

**Figure 5.**
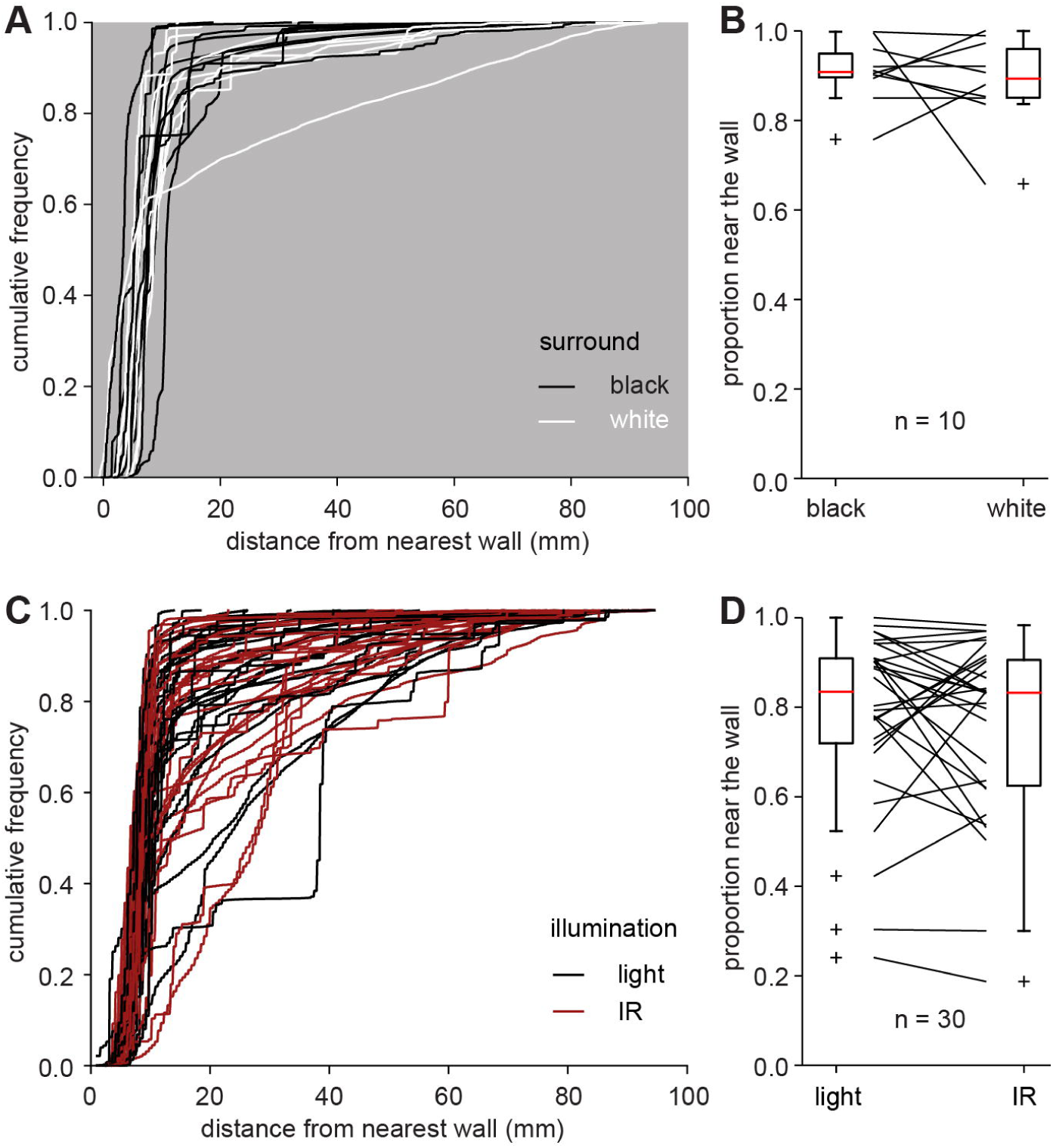
Influence of illumination conditions on wall following during swimming in *Xenopus* larvae. (A) Cumulative frequency distributions of the distance to the nearest wall during swimming of stage 53 – 65 tadpoles/froglets *(n* = 10) over a 10 min period in a 19 × 19 cm tank surrounded by black (black traces) or white paper (white traces). (B) Proportion of time that the animals spent near the wall (within 15 mm) for swimming in the tank surrounded by black (left) or white (right) paper. The proportions in these two conditions were not significantly different (paired t-test, *p* = 0.59). (C) Cumulative frequency distributions of the distance to the nearest wall during swimming of stage 50 – 65 tadpoles/froglets (*n* = 30) over a 10 min period in a 19 × 19 cm tank illuminated either with cold light (light, black traces) or infrared light (IR, red traces). (D) Proportion of the time that the animals spent near the wall (within 15 mm) for swimming in the tank with cold light (left) or IR light (right). The proportions in these two illumination conditions were not significantly different (paired Wilcoxon signed-rank test, *p* = 0.47).

### Influence of tank size on wall following

Wall following might be influenced by the size of the environment. To test whether the wall is equally attractive independent of the size of the tank, animals of developmental stages 47 – 50 (*n* = 9) were tested both in a 19 × 19 cm and in a 7 × 7 cm tank. The cumulative frequency distributions of distances to the nearest wall suggest that the animals spend more time near the wall in the smaller tank (Fig. 6A). This is confirmed by comparing the proportion of time that the animals spent near the wall (within 15 mm of the wall) in the two tanks: the proportions in the small tank are significantly larger (Fig. 6B, paired Wilcoxon signed rank test, *p* = 0.0078). This suggests that the wall is more attractive in the smaller tank. However, the ‘near wall’ area (within 15 mm of the wall) is also relatively larger in the smaller tank (67.3% of the total area in the 7 × 7 cm tank vs. 29.1% of the total area in the 19 × 19 cm tank). To compare wall following on the same scale, the distances to the wall were normalised to their maximum, and a threshold was chosen that resulted in an intermediate ‘near wall’ area (threshold of 28% of the maximal distance to the wall, resulting in a ‘near wall’ area of 48% of the total tank area; Fig. 6C). The proportion of time spent in these area-normalised ‘near wall’ areas was again significantly larger in the smaller tank (Fig. 6D, paired Wilcoxon signed rank test, *p* = 0.0078). The animals are therefore stronger wall followers in the smaller tank also when taking into account the differences in area.

**Figure 6.**
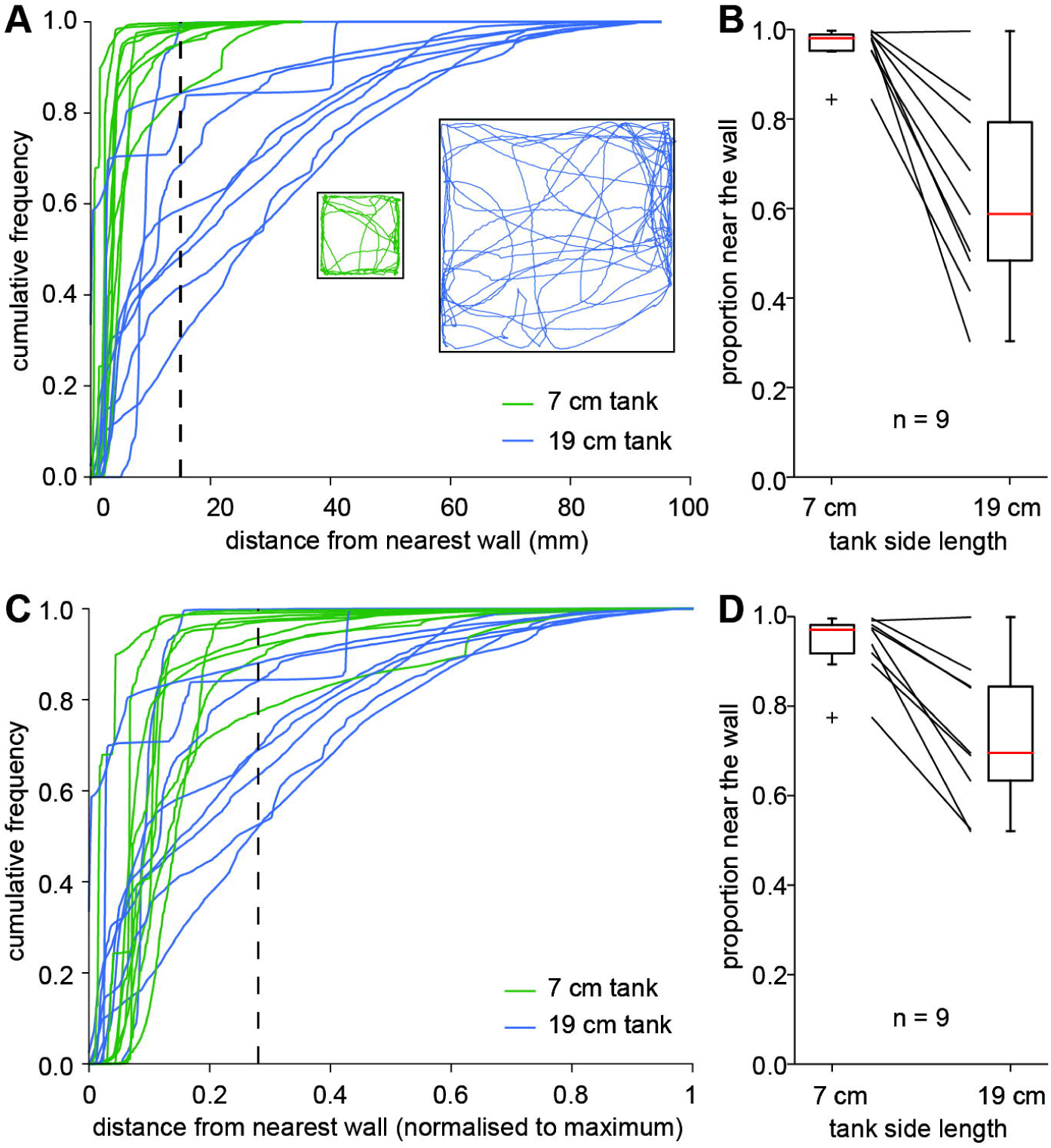
Influence of tank size on wall following. (A) Cumulative frequency distributions of the distance to the nearest wall during swimming of stage 47 – 50 tadpoles (*n* = 9) over a 10 min period in a 7 × 7 cm tank (green) and in a 19 × 19 cm tank (blue). The ‘near wall’ threshold (15 mm) is shown as a black dashed line. The trajectories of a stage 50 tadpole (3.2 cm body length) are shown as insets (B) Proportion of time that the animals in A spend near the wall (within 15 mm); the two groups were significantly different (paired Wilcoxon signed rank test, *p* = 0.0078, *n* = 9). (C) Same data as in A but normalised to the maximal distance to the wall. The black dashed line indicates the threshold (28% of the maximal distance to the wall) that yields a ‘near wall’ area intermediate to what 15 mm yields in the 7 and 19 cm tank (see Methods). (D) Proportion of time the animals spend near the wall (within 28% of the maximal distance) in the tanks with a side length of 7 and 19 cm; the two groups were significantly different (paired Wilcoxon signed rank test, *p* = 0.0078, *n* = 9).

### Wall following is passive

Wall following might be either active such as in blind cavefish (Patton et al., 2010) or passive (distinction according to Creed and Miller, 1990). To distinguish between the two possibilities for wall following in larval and adult *Xenopus*, the swimming behaviour was tested in a specifically designed tank (Fig. 7A,B). The use of a tank in which two of the four walls had convex curvatures allowed testing if tadpoles seek wall touch during swimming actively or follow concave walls passively (red and blue arrows in Fig. 7A). The proportion of trials when animals swam straight after encountering a convex curve (Fig. 7B) was evaluated from visual inspection by the experimenter. The majority of tested tadpoles swam straight in all trials (Fig. 7B) more or less independent of their developmental stage (Fig. 7C,D, *n* = 22), leading to the conclusion that wall following in *Xenopus* is passive.

**Figure 7.**
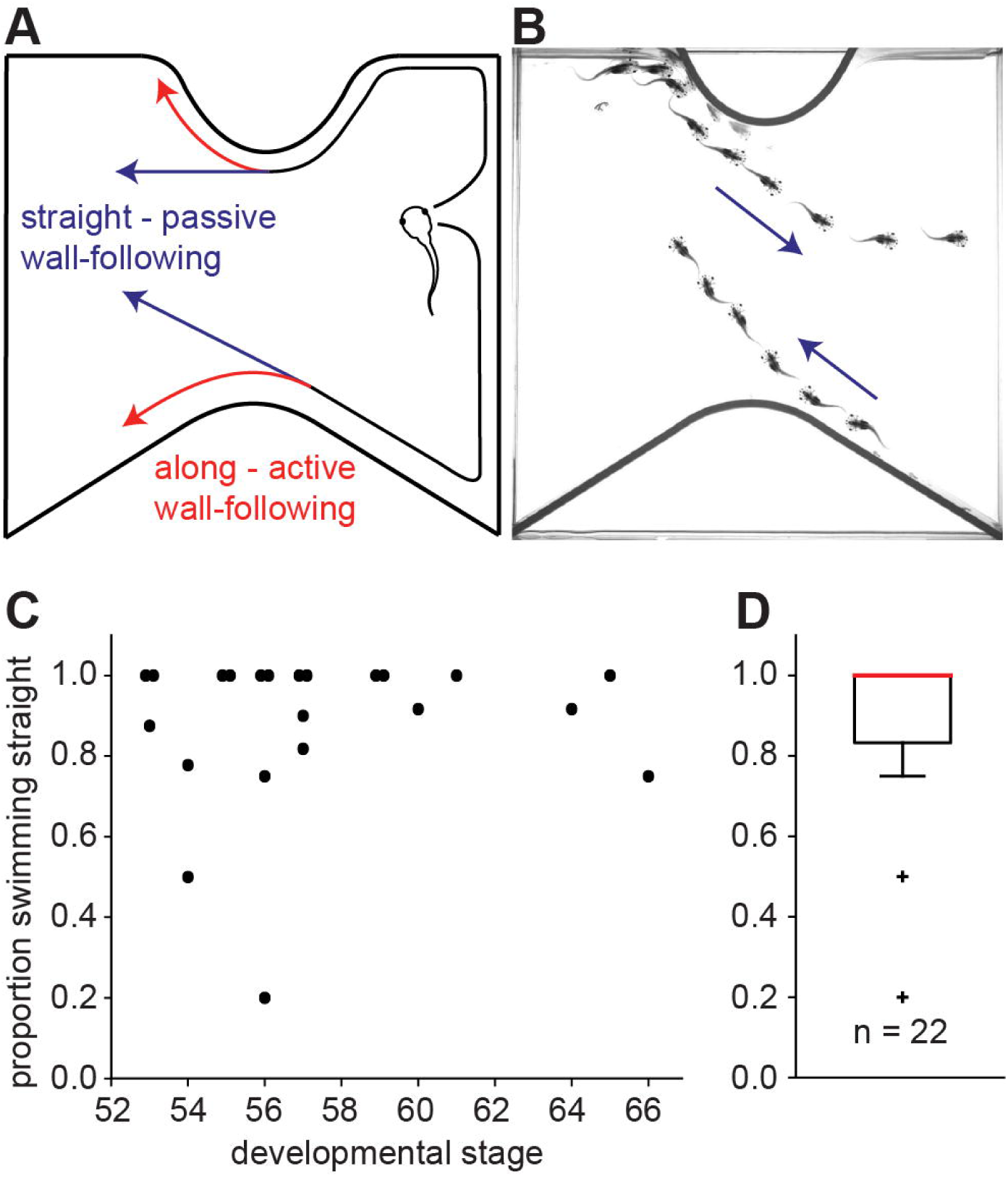
Wall following is passive. (A) Tank (19 × 19 cm) with two convex walls to distinguish if wall following is active (red arrows) or passive (blue arrows). (B) Minimum intensity overlay (at a frame rate of 3 fps) of two swimming trajectories along the curved walls of a stage 55 tadpole; blue arrows indicate the animal’s direction of swimming. (C) Proportion of trials with straight swimming and departure from the wall in animals at different developmental stages. (D) Boxplot of the proportion of straight swimming across all animals (*n* = 22). In C and D, only animals with at least 4 trials were included.

## Discussion

*Xenopus laevis* – from small tadpoles to froglets – tend to follow the wall when swimming in a square tank. The strength of wall following increases with progressive development and smaller tank size and is not confounded by the total distance that an animal covers. The transient presence of mechanosensory tentacles at mid-larval stages does not lead to stronger wall following compared to animals that naturally do not develop these appendages. Also, vision is unlikely a main driver of wall following, as surrounding the tank by black or white paper or changing the light to infrared illumination does not change the strength of wall following. Wall following is passive as indicated by straight swimming in a tank with convex curvatures. This indicates that wall following in *Xenopus* is likely imposed by the concave environment. Wall following being passive might also explain why it persists across metamorphosis and is present in both tadpoles and froglets, independent of their very different locomotor styles.

### Classification and different types of wall following

Wall following in concave environments has been described for a wide variety of animals: from crustaceans such as crayfish (Basil and Sandeman, 1999) to insects such as *Drosophila* (Besson and Martin, 2005; Martin, 2004), or cockroaches (Camhi and Johnson, 1999; Jeanson et al., 2003; Okada and Toh, 2000), to fishes such as zebrafish (Anichtchik et al., 2004; Colwill and Creton, 2011), goldfish (Kato et al., 1996), salmon (Clements et al., 2002) or blind cavefish (Patton et al., 2010; Teyke, 1985; Teyke, 1989), to several rodent species including voles, rats and mice (Eilam, 2004; Perrot-Sinal et al., 1999; Simon et al., 1994; Treit and Fundytus, 1988; Webster et al., 1979; Wilson et al., 1976). In many cases, these examples of wall following behaviours have been described in the context of thigmotaxis and centrophobism, and in relation to the level of anxiety. Thigmotaxis is a term that describes the motion of an organism relative to a touch stimulus; it is often used as shorthand for positive thigmotaxis, which means that animals actively seek out touch stimulation as they move. Centrophobism, on the other hand, is a tendency of animals to avoid open spaces, for instance the centre of an open test field for mice or rats (Martínez et al., 2002). Some authors use the term centrophobism when the avoidance of open spaces is related to vision (Cardenas et al., 2001). For instance common spiny mice move much more often into an open space in the dark than in the light (Eilam, 2004), though some authors use the term centrophobism without necessarily implying a visual mechanism. Thus, centrophobism and thigmotaxis are two potential but not mutually exclusive mechanisms that can lead to the avoidance of open spaces and the following of environmental boundaries. Wall following is therefore a neutral term to describe the tendency of an animal to follow vertical walls in its environment without a reference to the underlying mechanism. An environment with convex borders allows distinguishing between passive and active wall following (Creed and Miller, 1990). Animals perform active wall following when voluntarily seeking out the proximity to a wall and turn in order to remain near the wall. Passive wall following occurs when animals leave the wall at a convex curve but follow the walls in a concave environment. When wall following is active, thigmotaxis, centrophobism or a combination of the two can be the underlying mechanism.

### Potential uses of wall following

Thigmotaxis has been described both as a defensive strategy (Grossen and Kelley, 1972) as well as a spatial exploration strategy (Kallai et al., 2007). Animals might be safer near a vertical wall compared to the open; for instance it has been suggested that avian predation on rats likely is lower near a wall than in the open (Grossen and Kelley, 1972). Mice increase thigmotaxis in the presence of a potential predator (Bonsignore et al., 2008). Other rodents such as the common spiny mouse only venture in the centre of an open field if there are objects that might serve as shelter, or if it is dark (Eilam, 2004). Moreover, thigmotaxis has been related to anxiety, and is commonly used as a simple behavioural readout of anxiety levels in mice and rats (Prut and Belzung, 2003; Simon et al., 1994; Treit and Fundytus, 1988). Some authors argue that fear of open spaces is not only driven by touch but also by vision (Martínez et al., 2002), which suggests that wall hugging and avoidance of open spaces is a combination of thigmotaxis and centrophobism. Independent of the underlying mechanisms, the use as a defensive strategy is clear. Moreover, wall following can also serve as a useful spatial exploration strategy. Especially under conditions when long-range sensing such as vision is not available, exploration of the environment based on touch along its borders can provide the basis for the formation of a cognitive map (Kallai et al., 2007; Yaski et al., 2009) and serve as a reference frame for later exploration (Kallai et al., 2005). However, this is only useful as an initial strategy; if it is used excessively it can even prevent further spatial learning (Kallai et al., 2007). Such initial wall following as a means for spatial learning has been observed in various species such as crayfish (Basil and Sandeman, 1999), blind cavefish (Teyke, 1989), and blind mole rats (Avni et al., 2008).

### Persistence of wall following with development in Xenopus

In this study we examined a range of developmental stages of *Xenopus* – from small to large tadpoles immediately prior to metamorphosis as well as froglets after metamorphosis has been completed. Wall following in a square tank was present at all developmental stages; the strength of wall following was weakest, however, in the smallest tadpoles, stronger, with considerable variations in larger tadpoles and consistently strong in froglets. This persistence suggests that wall following is not a behavioural strategy only employed by tadpoles or frogs, and is not linked to a particular locomotor style such as undulatory tail-based propulsion or leg-based swimming. Moreover, wall following in a convex tank was passive in all animals tested (see below). The weaker wall following in young larvae is noticeable and might be related to the somewhat different swimming style of these animals (see Fig. 3A in Hänzi and Straka, 2017), where the rotation axis of the left-right head undulations oscillates between positions outside the animal; this is at variance with the situation in larger tadpoles where the head oscillations during swimming occur around a single central axis (Lambert et al., 2009). This difference in swimming style might facilitate turns away from a vertical wall in young larvae and explain the weaker wall following.

At intermediate developmental stages examined in this study (stage 51-60 according to Nieuwkoop and Faber, (1956)), tadpoles normally possess a pair of mobile appendages protruding from the corners of their mouths, which are retracted during undulatory swimming (Hänzi et al., 2015). These tentacles – like other skin areas – possess mechanoreceptive Merkel cells (Nurse et al., 1983; Ovalle, 1979; Ovalle et al., 1998), and therefore the tentacles likely serve a tactile function when the animal is stationary or cruising slowly with tentacles extended forward. We hypothesised that these tentacles might be used to explore the environment in a way that is similar to rodents’ whiskers but simpler because the structure is not as specialised. However, younger larvae and older animals at metamorphic climax (>stage 61) that do not possess any tentacles were overall similar in their wall following tendencies, as were animals that for unknown reasons did not develop tentacles (Fig. 4). While this does not exclude that – when present – tentacles are used for tactile exploration, it shows at least that tentacles are not necessary for wall following, and if tactile exploration is needed, tadpoles might also use their facial skin.

### Effects of vision

As mentioned above, some rodents leave the walls and venture much more into open space in darkness than in light; this is true not only for the common spiny mouse (Eilam, 2004) but also for rats (Nasello et al., 1998), some types of gerbil (Zadicario et al., 2005) and wild-caught prairie deer mice (Brillahart and Kaufman, 1991). Some rodents also adjust their foraging behaviour in laboratory or natural conditions such that they venture more into the open in the dark (Diaz, 1992; Price et al., 1984; Vasquez, 1996), and some authors also assign a role of vision in the avoidance of open spaces by rats (Cardenas et al., 2001; Martínez et al., 2002). However, tadpoles and froglets of *Xenopus laevis* did not show stronger wall following in light than under infrared illumination. While the IR lamps used here did not produce pure infrared light, IR illumination nevertheless is a condition with considerably reduced light and influence of vision. Centrophobism or visually driven fear of open spaces is therefore very unlikely to be the driving force behind wall following in *Xenopus*. The wall following strategy might rather be a side effect of locomotion in the mostly murky aquatic environment of the natural habitat of *Xenopus* (Nieuwkoop and Faber, 1956) independent of the developmental stage.

### Effects of the size of the environment

A range of different arena sizes have been used in rodent open field tests (Walsh and Cummins, 1976), and the geometry of the environment has shown to influence path shapes of rats not only at the perimeter but also at the centre of an environment (Yaski et al., 2011). A wall can exert both a guiding and attracting influence on mouse behaviour from quite some distance (Horev et al., 2007). Two studies explicitly examined the proportion of time that social voles spend near the wall in arenas of different sizes (Eilam, 2003; Eilam et al., 2003). These animals are very active, and spend more time near the wall in larger arenas – possibly because the larger open space is perceived as more dangerous than a smaller, more enclosed open space. This contrasts with the behaviour of *Xenopus* described here, which are stronger wall followers in smaller tanks. It therefore seems likely that wall following in *Xenopus* is imposed by the constraints of the environment, whereas wall following in social voles serves as a defensive strategy. Moreover, thigmotaxis is unlikely to be the main mechanism behind wall following in either of the two cases, since different tank/arena sizes would have no impact on wall following if thigmotaxis was the underlying cause (Eilam et al., 2003) – with thigmotaxis as the main mechanism, the walls would be equally attractive independent of the arena size.

### Passive versus active wall following

The studies ascribing protection or exploration as the function of wall following have used the terms thigmotaxis or centrophobism for a reason: wall following can only be protective or exploratory if it is active. Passive wall following such as observed in this study in *Xenopus laevis* is rather unlikely to serve these purposes. To the best of our knowledge, no other study described passive wall following so far. Potential reasons include that only few studies use convex tanks, and that passive wall following might be considered a negative finding and not be reported. The few following studies did use convex enclosures to discriminate active from passive wall following: In blind cavefish, for instance, wall following is clearly active (Patton et al., 2010; Sharma et al., 2009). These animals are blind, live in dark caves and use their lateral line system as a near range sense to obtain information about their environment. In a convex tank they actively follow the wall because they would not be able to orient otherwise. Adult fruit flies, on the other hand, leave the wall in more than 50% of the trials; their preference for walls in circular arenas seems to derive from a preference for the boundaries of the environment rather than from thigmotaxis or centrophobism (Soibam et al., 2012). In contrast to fruit flies, cockroaches have antennae that can be longer than their body (Camhi and Johnson, 1999). These animals use these mechanoreceptive sensors to gain information about their nearby environment. Cockroaches thus have been described as thigmotactic in concave environments (Camhi and Johnson, 1999; Jeanson et al., 2003) and show positive thigmotaxis towards objects that are touched by the antennae (Okada and Toh, 2000). When running along a wall these animals constantly touch the wall with one of their antennae (Camhi and Johnson, 1999). However, when arriving at a convex curve, they leave the curve in about 50% of the trials (Creed and Miller, 1990).

Active wall following in blind cavefish certainly serves as a spatial exploration and spatial learning strategy (Teyke, 1989), and to a certain extent this might also be true for cockroaches or fruit flies. In contrast, wall following in *Xenopus laevis* is passive and therefore unlikely to serve as a specific protective or exploratory strategy or a behaviour that is related to anxiety. A number of factors potentially influencing wall following such as changes in illumination or the presence of tentacles were shown to play no major role for wall following in *Xenopus*. Instead, passive wall following in these animals might be due to the particularity of the rather unnatural and concave test environment. This thus suggests that spatially more complex and natural environments likely would yield richer behaviours (see also Benjamini et al., 2010; Cheng, 2005).

**List of abbreviations:** IR: infrared.

## Acknowledgements

The authors thank all members of the Straka lab for feedback and discussion.

**Competing interests** No competing interests are declared.

**Author contributions**

Investigation, Software, Visualization: S.H.; Supervision, Project administration, Funding acquisition: H.S.; Conceptualization, Writing: S.H. and H.S.

